# Separable, Ctf4-mediated recruitment of DNA Polymerase α for initiation of DNA synthesis at replication origins and lagging-strand priming during replication elongation

**DOI:** 10.1101/352567

**Authors:** Sarina Y. Porcella, Natasha C. Koussa, Colin P. Tang, Daphne N. Kramer, Priyanka Srivastava, Duncan J. Smith

**Affiliations:** Department of Biology, New York University, New York, NY 10003, USA; Weill-Cornell Graduate School of Medical Sciences, New York, NY 10065, USA; School of Dentistry, UCSF Center for the Health Sciences, San Francisco, CA 94143

**Keywords:** DNA replication, Lagging-strand synthesis, Replication origins, Checkpoint activation

## Abstract

During eukaryotic DNA replication, DNA polymerase alpha/primase (Pol α) initiates synthesis on both the leading and lagging strands. It is unknown whether leading- and lagging-strand priming are mechanistically identical, and whether Pol α associates processively or distributively with the replisome. Here, we titrate cellular levels of Pol α in *S. cerevisiae* and analyze Okazaki fragments to study both replication initiation and ongoing lagging-strand synthesis *in vivo*. We observe that both Okazaki fragment initiation and the productive firing of replication origins are sensitive to Pol α abundance, and that both processes are disrupted at similar Pol α concentrations. When the replisome adaptor protein Ctf4 is absent or cannot interact with Pol α, lagging-strand initiation is impaired at Pol α concentrations that still support normal origin firing. Additionally, we observe that activation of the checkpoint becomes essential for viability upon severe depletion of Pol α. Using strains in which the Pol α-Ctf4 interaction is disrupted, we demonstrate that this checkpoint requirement is not solely caused by reduced lagging-strand priming. Our results suggest that Pol α recruitment for replication initiation and ongoing lagging-strand priming are distinctly sensitive to the presence of Ctf4. We propose that the global changes we observe in Okazaki fragment length and origin firing efficiency are consistent with distributive association of Pol α at the replication fork, at least when Pol α is limiting.

**Author summary:** Half of each eukaryotic genome is replicated continuously as the leading strand, while the other half is synthesized discontinuously as Okazaki fragments on the lagging strand. The bulk of DNA replication is completed by DNA polymerases ε and δ on the leading and lagging strand respectively, while synthesis on each strand is initiated by DNA polymerase α-primase (Pol α). Using the model eukaryote *S. cerevisiae*, we modulate cellular levels of Pol α and interrogate the impact of this perturbation on both replication initiation on DNA synthesis and cellular viability. We observe that Pol α can associate dynamically at the replication fork for initiation on both strands. Although the initiation of both strands is widely thought to be mechanistically similar, we determine that Ctf4, a hub that connects proteins to the replication fork, stimulates lagging-strand priming to a greater extent than leading-strand initiation. We also find that decreased leading-strand initiation results in a checkpoint response that is necessary for viability when Pol α is limiting. Because the DNA replication machinery is highly conserved from budding yeast to humans, this research provides insights into how DNA replication is accomplished throughout eukaryotes.

## Introduction

DNA polymerase alpha/primase (Pol α) is responsible for initiating synthesis on the leading strand and for each Okazaki fragment on the lagging strand (Kunkel and Burgers, 2008). The ultimate contribution of Pol α to replication is limited, and bulk synthesis on the leading- and lagging-strands is carried out by DNA polymerase epsilon (Pol ε) and polymerase delta (Pol δ), respectively (Clausen et al., 2015; Daigaku et al., 2015; Pursell et al., 2007; Reijns et al., 2015). Despite the fact that different DNA polymerases carry out the majority of DNA synthesis on the two daughter strands, Pol α hands off synthesis predominantly to Pol δ during normal initiation on both strands and during replication restart (Daigaku et al., 2015; Garbacz et al., 2018; Yeeles et al., 2017; Miyabe et al., 2015). The use of the same initiating polymerase and downstream partner implies a possible similarity between leading- and lagging-strand initiation. Indeed, recent evidence from reconstituted replisomes suggests that leading-strand initiation can occur via extension of the first Okazaki fragment synthesized on the lagging strand (Aria and Yeeles, 2018). However, analysis of repriming during damage bypass in the same reconstituted system implies that leading- and lagging-strand repriming may be mechanistically distinct (Taylor and Yeeles, 2018).

Eukaryotic Okazaki fragments are considerably shorter than their prokaryotic counterparts (Balakrishnan and Bambara, 2013; Ogawa and Okazaki, 1980). Okazaki fragment length is quantized by the nucleosome repeat via interactions between nascent nucleosomes and Pol δ in *S. cerevisiae* (Smith and Whitehouse, 2012) and *C. elegans* (Pourkarimi et al., 2016). Okazaki fragment termini can also be positioned by nucleosomes in a reconstituted *S. cerevisiae* replication reaction (Devbhandari et al., 2017). Both the distribution of Okazaki fragment termini with respect to nucleosomes and the overall length profile of lagging-strand products in *S. cerevisiae* and *C. elegans* are remarkably similar. Despite this apparent size conservation, Okazaki fragment length can be altered on naked and non-chromatinized templates *in vitro* by varying the concentration of Pol α (Kurat et al., 2017), analogous to the impact of primase titration on lagging-strand synthesis in a reconstituted *E. coli* replication system (Wu et al., 1992). Eukaryotic Okazaki fragment length can also be increased *in vivo* by impairing nucleosome assembly (Smith and Whitehouse, 2012; Yadav and Whitehouse, 2016).

The maximum length of an Okazaki fragment is determined by the amount of single-stranded DNA unwound at the replication fork before lagging-strand priming and extension. Thus, longer fragments would result in the exposure of long stretches of damage-prone single-stranded DNA. Shorter Okazaki fragments would expose shorter stretches of ssDNA, but at the likely cost of increasing the contribution of the error-prone Pol α to synthesis of the lagging daughter strand (Kunkel, 2011; Reijns et al., 2015). However, it is currently unclear to what extent changing Okazaki fragment length directly impacts cellular fitness. Reduced DNA polymerase activity upon aphidicolin treatment induces chromosome breakage and genomic rearrangements at fragile sites in mammals (Glover et al., 1984). Analogously, reducing the intracellular concentration of Pol α in *S. cerevisiae* increases S-phase duration, sensitivity to DNA damaging agents, and chromosomal rearrangements at defined sites (Lemoine et al., 2005; Song et al., 2014). The detrimental effects of Pol α depletion on genome integrity could arise due to defective leading- or lagging-strand initiation, both of which are dependent on Pol α.

Ctf4 was originally identified as a chromosome transmission fidelity mutant (Kouprina et al., 1992). Subsequent studies have identified multiple roles for Ctf4 in DNA metabolism. Ctf4 is not only required for the establishment of sister chromatin cohesion (Borges et al., 2013; Hanna et al., 2001), but also links Pol α to the replication fork via interaction with the replicative helicase (Gambus et al., 2009; Tanaka et al., 2009), and is required for error-free lesion bypass (Fumasoni et al., 2015) and rDNA maintenance (Villa et al., 2016). At least some of these roles for Ctf4 are independent of Pol α binding. The metazoan Ctf4 ortholog, AND-1 also stimulates Pol α binding to chromatin, and is required for efficient DNA replication (Zhu et al., 2007). Interestingly, despite the conserved contribution of Ctf4 to Pol α recruitment, *ctf4Δ S. cerevisiae* strains have been demonstrated to synthesize identically sized Okazaki fragments to wild-type cells (Borges et al., 2013). Additionally, the Ctf4 protein has a minimal effect on the priming of either DNA strand in reconstituted replication reactions (Kurat et al., 2017). Thus, the contributions of Ctf4 to Pol ⍺ recruitment for origin firing and Okazaki fragment initiation have not been fully elucidated *in vivo*.

Here, we analyze Okazaki fragments to directly test the effects of Pol α depletion on origin firing efficiency and Okazaki fragment initiation in *S. cerevisiae*. We find that reduced levels of Pol α lead to an increase in Okazaki fragment length and a global decrease in replication-origin firing efficiency. In the absence of a Ctf4-Pol1 interaction, lagging-strand initiation is impaired at moderate Pol α concentrations that support normal levels of origin firing. Impaired Okazaki fragment initiation is well tolerated: however, a severe reduction in Pol α levels leads to a strict dependence on checkpoint activation for continued viability.

## Results

### Cells with reduced levels of Pol α synthesize longer Okazaki fragments *in vivo*

To investigate the effect of Pol1 depletion on leading- and lagging-strand priming during replication, we modified the approach of Petes and co-workers (Lemoine et al., 2005; Song et al., 2014), limiting expression of *POL1* by replacing its promoter with *pGAL1*. Expression of Pol1 from galactose-inducible promoters generates a stable polypeptide that can persist for several cell cycles (Muzi Falconi et al., 1993). Therefore, to facilitate turnover of pre-existing Pol1 and focus on the acute effects of depletion, we additionally fused an N-terminal degron to the *POL1* coding sequence. Western-blot analysis indicated that the concentration of this GAL1-expressed, degron-tagged Pol1 (GDPol1) could be specifically and rapidly modulated within 4h (Fig. 1A). The lower band marked with an asterisk in GDPol1-myc Western blots is a degradation product resulting from degron-tagging (see Fig. S1A). All cultures were grown in media supplemented with 3% raffinose to avoid indirect effects due to carbon limitation. Pol1 levels oscillate through the cell cycle when the protein is endogenously expressed, but not when expression is driven by galactose (Muzi Falconi et al., 1993). Because Pol1 concentration is not constant even within S-phase (Muzi Falconi et al., 1993), and only 2/3 of cells in an asynchronous population are in G1, G2 or M phase, we compared wild-type POL1 expression during S phase to 0.5% and 0.05% galactose in our inducible system at the zero and 60 minute timepoints after release from alpha-factor-mediated G1 arrest (Fig. S1A). We estimate that the expression level at 0.05% galactose is slightly lower than endogenous during the relevant phase of the cell cycle, consistent with several phenotypes described below. However, we note that the concentration of free Pol ⍺ might vary through the cell cycle as the complex associates with elongating replication forks, especially under limiting conditions.

**Figure 1.**
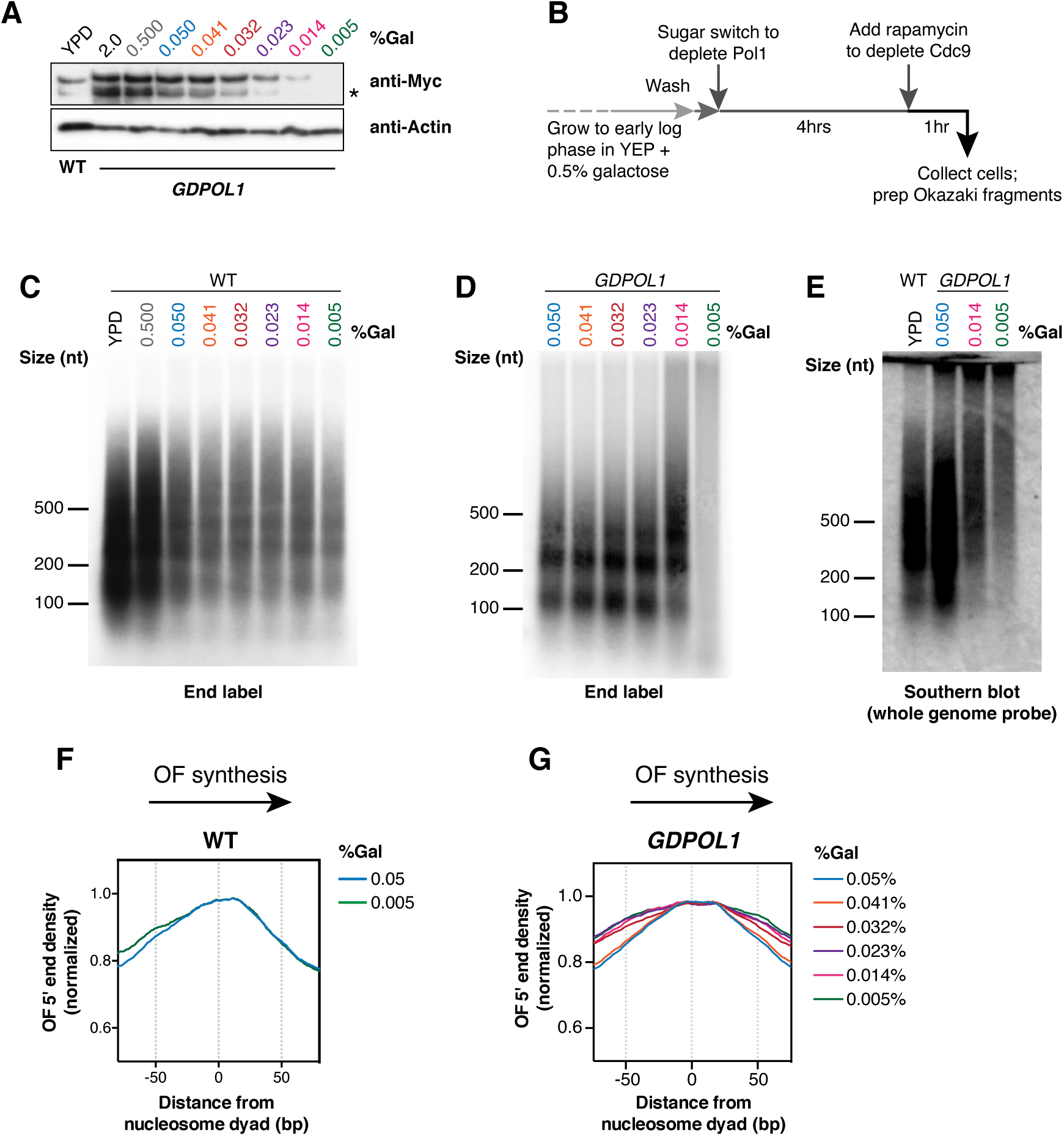
Okazaki fragments increase in length when Pol α is limiting. **(A).** Western blot against 13xMyc-tagged Pol1 from S. cerevisiae from a wild-type or *GDPOL1* strain, as indicated, shifted to YPD or YEP + 3% raffinose (hereafter, media) supplemented with various concentrations of galactose. The lower band indicated by an asterisk is a degradation product resulting from degron-tagging. **(B).** Schematic of experimental workflow for Okazaki fragment analysis and sequencing (also see methods). **(C, D).** Alkaline agarose gel analysis of end-labeled Okazaki fragments from a wild type (C) or *GDPOL1* (D) strain, shifted to YPD or media supplemented with galactose as indicated. **(E).** Southern blot using a whole genome probe, on Okazaki fragments from a wild type or *GDPOL1* strain shifted to media with the indicated sugar concentrations. **(F, G).** Distribution of Okazaki fragment 5’ ends around consensus nucleosome dyads (Jiang and Pugh, 2009) for a wild-type (F) or *GDPOL1* (G) strain shifted to media containing the indicated concentration of galactose. Data for Okazaki fragment 3’ ends in *GDPOL1* are in Fig. S1F

To analyze Okazaki fragment biogenesis, we crossed the *GDPOL1* allele into a strain background in which DNA ligase I (Cdc9) can be depleted from the nucleus by rapamycin treatment using the anchor away method (Haruki et al., 2008). Nuclear depletion of Cdc9 enriches nucleosome-sized Okazaki fragments (Fig. S1B), similarly to transcriptional repression of Cdc9 (Smith and Whitehouse, 2012; Yadav and Whitehouse, 2016). Robust detection of Okazaki fragments was possible after 1h rapamycin treatment: therefore, all Okazaki fragment labeling and sequencing experiments were conducted after a 4h sugar switch to reduce Pol α levels, followed by 1h ligase depletion by rapamycin (Fig. 1B).

By end-labeling unligated Okazaki fragments, we observed that cells with wild-type *POL1* did not show an increase in Okazaki fragment length under low-galactose growth conditions (Fig. 1C). This result was highly reproducible (another representative gel is shown in Fig. S1C). By contrast, in the *GDPOL1* strain Okazaki fragment length was normal at Pol1 concentrations down to a critical concentration corresponding to growth in 0.014% galactose (Fig. 1D, representative replicate experiments are shown in Fig. S1D-E). At 0.014% galactose the Okazaki fragment length profile was shifted slightly upwards such that fragments were clearly still phased by nucleosomes, while lower galactose concentrations (0.005%) showed a significant loss of signal (Fig. 1D cf. lanes 5&6). To confirm that this loss of signal reflected a further length increase (and therefore a reduction in the number of ends being labeled), we analyzed Okazaki fragments by Southern blot using a whole-genome probe: as anticipated for severely perturbed lagging-strand priming, Okazaki fragments at 0.005% galactose were significantly larger than at 0.014% (Fig. 1E). We conclude that limiting levels of Pol α lead to reduced priming frequency on the lagging strand, resulting in longer Okazaki fragments, and that *S. cerevisiae* cells can sustain growth when Okazaki fragment length is substantially increased. These data are consistent with the presence of multiple Pol ⍺ complexes at the replication fork and/or the repeated, distributive recruitment of Pol α to the replisome for lagging-strand priming during replication, since the priming kinetics of a single Pol α complex stably associated with the replisome would be unaffected by cellular Pol α concentrations (see discussion).

To analyze the location of Okazaki fragment termini, we purified and sequenced Okazaki fragments (Smith and Whitehouse, 2012) from wild-type and *GDPOL1* strains shifted to low galactose concentrations for 4h before 1h ligase depletion. Galactose concentration does not significantly affect the distribution of Okazaki fragment termini in wild-type cells (Fig. 1F). However, we observed that both the 5’ (Fig. 1G) and 3’ (Fig. S1F) termini of Okazaki fragments in *GDPOL1* cells were less enriched at nucleosome dyads during growth at concentrations below 0.041% galactose (Fig. 1G). Pol1 interacts with the FACT component Spt16 (Foltman et al., 2013): furthermore, Pol α contains a histone-binding motif for H2A and H2B and it is implicated in the maintenance of repressive chromatin during replication (Evrin et al., 2018). The change in the distribution of Okazaki fragment ends at moderate Pol α levels that do not affect lagging-strand priming (cf. Fig. 1D&G) supports an intimate role for Pol ⍺ in chromatin assembly on the lagging strand.

### Reduced levels of Pol α lead to a global decrease in replication origin firing efficiency

Replication origin firing efficiency can be quantitatively inferred from the distribution of Watson- and Crick-strand Okazaki fragments after deep sequencing (McGuffee et al., 2013; Petryk et al., 2016; Pourkarimi et al., 2016). By comparing the fraction of Okazaki fragments mapping to the Watson and Crick strands in the region ±10 kb from the replication origin midpoint, an Origin Efficiency Metric (OEM) can be calculated (McGuffee et al., 2013). Okazaki fragment distributions across a ∼400 kb region of chromosome 4 containing both early and late firing regions are shown in Fig. 2A. Genome-wide origin efficiencies are quantified as OEM in Fig. 2B. OEMs for each origin at all conditions are compiled in Table S1. Comparisons between replicate datasets were robust across galactose concentrations (Fig. S2A), and each Pol α concentration maintains a consistent trend across replicates (Fig. S2B). Data in Fig. 2 represent the mean origin efficiency across three replicate experiments. In wild-type cells, origin efficiency was unaffected across the full range of galactose concentrations tested (Fig. 2B). In the *GDPOL1* strain, origin firing was maintained at wild-type levels above a critically low level of Pol α (0.023% galactose). A significant reduction in average origin firing efficiency was observed at 0.023% galactose, and firing efficiency was maintained at this decreased level at progressively lower galactose concentrations (Fig. 2B). These changes in average origin firing efficiency are also shown in Fig. 2C as a change in the proportion of Okazaki fragments mapping to the Watson strand around a meta-origin.

**Figure 2.**
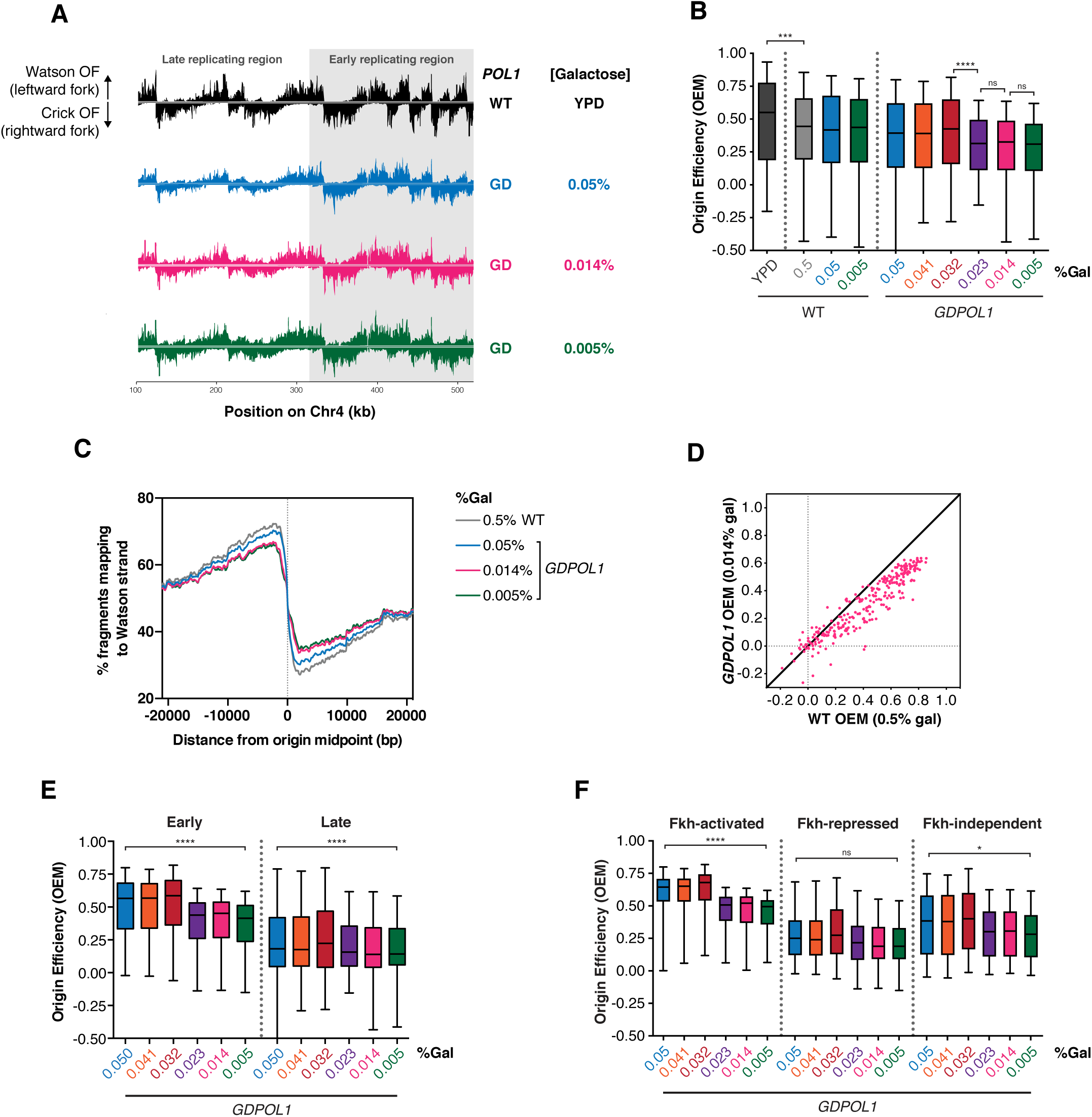
Origin firing efficiency is decreased for all replication origins when Pol α levels are reduced. **(A).** Okazaki fragment distributions from a wild-type (black) or *GDPOL1* strain shifted to the indicated media. A ∼400 kb region from the left arm of chromosome 4 is shown, and the late and early replicating regions are annotated. **(B).** Origin efficiency calculated as OEM (see methods) in wild-type or *GDPOL1* cells shifted to the indicated concentration of galactose. Data is in the form of box and whisker plots with the ends of the boxes being the upper and lower quartiles, the median is denoted by the line within the box, and the whiskers indicate the highest and lowest data points. Origin efficiency was calculated as in (McGuffee et al., 2013), using origin locations annotated in the same study. Significance was calculated by unpaired t-test; **** p<0.0001, *** p<0.0005, * p<0.05. Data from the *GDPOL1* strain are the average of three replicates. **(C).** Meta-analysis of the fraction of Okazaki fragments mapping to the Watson strand (corresponding to leftward-moving replication forks) around all 283 origins normalized to the maximum, using the same origin list as above. **(D).** Scatter plot comparing origin firing efficiency in *GDPOL1* cells at 0.014% galactose, to wild-type cells in 0.5% galactose. Analogous comparisons to other galactose concentrations, and correlations between replicates, are in Fig. S2. **(E).** Firing efficiency for origins with replication timing below (early) or above (late) the median replication timing for origins in our dataset. Significance was calculated by unpaired t-test; **** p<0.0001. **(F).** Firing efficiency for origins with Forkhead (Fkh) status: activated, repressed, or independent. Using the Fkh status determined in Knott et al., 2012. Significance was calculated by unpaired t-test; **** p<0.0001, * p<0.05.

Interestingly, we observed that the efficiency of productive replication origin firing is impaired at a similar Pol1 concentration (0.023% galactose) to the concentration at which Okazaki fragment length is increased (0.014% galactose) (Fig 1D&E). This decrease in origin efficiency likely contributes to the slow growth (Fig. S3A), cell-cycle delay (Fig. S3B & (Lemoine et al., 2005)), and accumulation of S-phase cells (Fig. S3C) observed under limiting Pol1 conditions. However, since growth is slower at 0.005% than 0.014% while origin efficiency is unchanged, reduced origin firing cannot be the only cause of the slow growth. To test whether replisome stalling or arrest at hard-to-replicate sites was increased under low Pol1 conditions, we analyzed replication direction around 93 tRNA genes (Fig. S4) as previously described (Osmundson et al., 2017). tRNA genes are the major sites of replisome stalling in the *S. cerevisiae* genome (Ivessa et al., 2003; Osmundson et al., 2017), and changes in Okazaki fragment polarity can robustly detect increased or decreased fork stalling at these sites (Osmundson et al., 2017). We did not observe fork stalling at any galactose concentration (Fig. S4). Thus, we conclude that replication-fork stalling or arrest is not substantially affected by limiting Pol ⍺ concentrations.

If one or more Pol α complexes is stably recruited to and maintained in the replisome upon leading-strand initiation, reducing Pol α to sub-stoichiometric levels would privilege early and/or efficient replication origins while disproportionately reducing the efficiency of late and/or inefficient origins. By contrast, distributive recruitment would lead to reduced efficiency of all origins under limiting Pol α conditions. We analyzed origin efficiency in the *GDPOL1* strain as a function of normal firing efficiency. As the concentration of Pol α was decreased below the threshold, firing of essentially all origins became less efficient regardless of their normal efficiency (Fig. 2D & S2C). Both early- and late-firing replication origins showed a global decrease in firing efficiency at low Pol1 concentrations (Fig. 2E). Similarly, origins whose firing is stimulated or unaffected by forkhead-mediated spatial clustering (Knott et al., 2012) were all affected by Pol ⍺ depletion (Fig. 2F). The firing efficiency of origins repressed by forkhead transcription factors was not significantly impacted by Pol ⍺ depletion (Fig. 2F), likely because these origins fire inefficiently under normal conditions.

To confirm the global decrease in origin firing efficiency at limiting Pol α levels, we analyzed DNA copy number via whole genome sequencing (WGS) of cells collected in S-phase following release from G1 arrest at a range of galactose concentrations (Fig. 3A). We sequenced samples from early, mid, and late S-phase cells based on flow cytometry grown at 0.05%, 0.014% and 0.005% galactose (Fig. S3D), and combined these data to obtain a snapshot of the population across the whole of S-phase.

**Figure 3.**
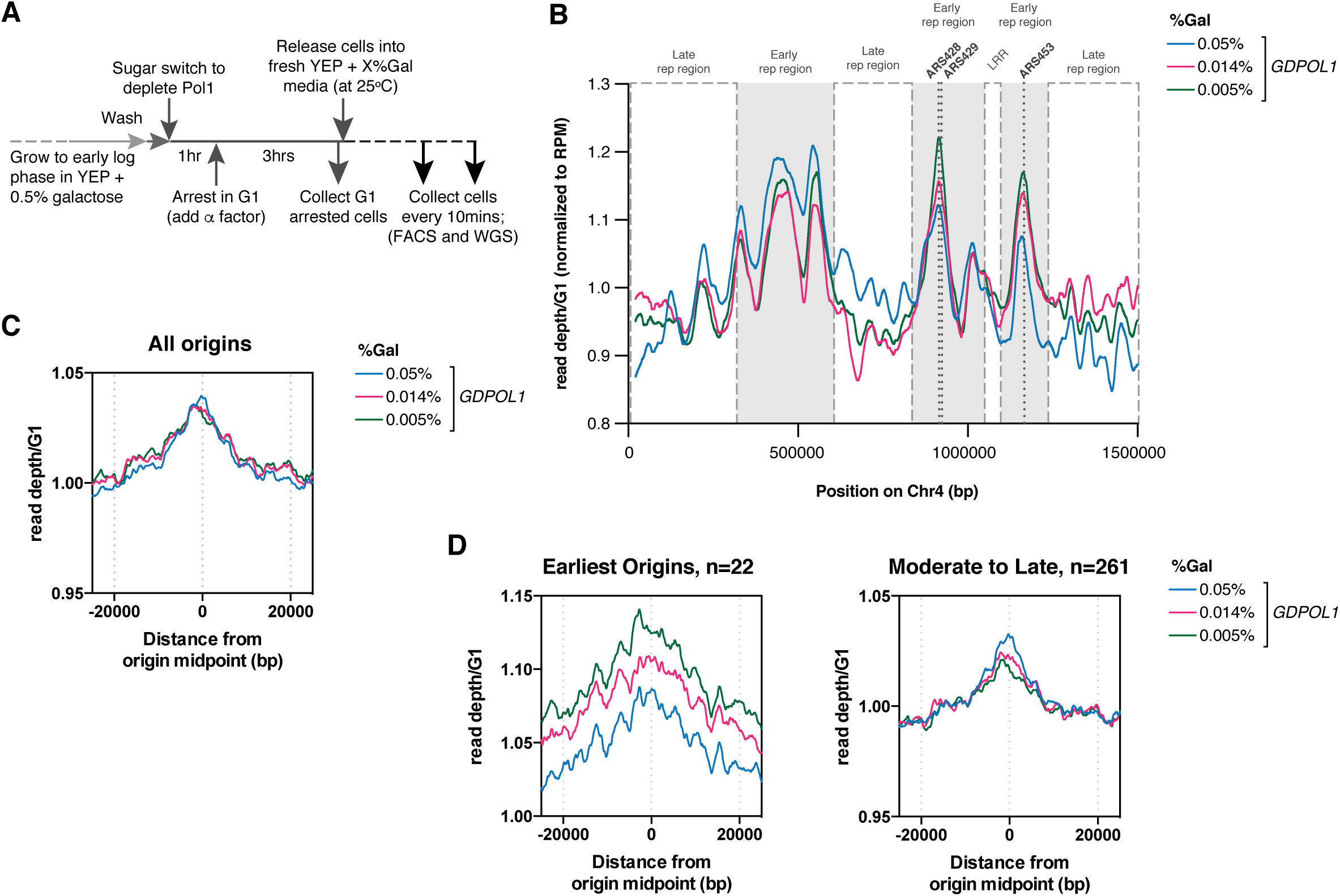
Global decreases in origins firing from Pol α depletion leads to a proportion of cells stalling in early S phase. **(A).** Schematic of experimental workflow for whole genome sequencing (also see methods). **(B)** Read depth of chromosome 4, normalized to G1 and RPM, then smoothed through a Loess regression. Poorly mapped regions were removed for analysis (Table S2). Galactose concentrations are indicated. Early and late replication regions (LRR) are marked, as are the three earliest firing origins on chromosome 4: ARS428 (12min), ARS429 (14min), ARS453 (12min). **(C, D).** Coverage from pan-S-phase samples mapped around replication origins. Either all 283 origins (C) or the earliest firing 22 origins compared to the other 261 origins (D) are shown. Read depth was normalized to G1 and smoothed to 1kb. Note the different y-axis scale for the earliest origins in D.

The read depth across chromosome 4 from our pan-S-phase samples, normalized to a G1 sample, is shown in Fig. 3B. The global reduction in origin firing efficiency inferred from Okazaki fragment sequencing (Fig. 2A-D) would be expected to ‘flatten’ the distribution of read depths in S-phase such that early-replicating regions are less overrepresented and late-replicating regions are correspondingly less underrepresented. Our data are broadly consistent with this prediction (cf. labeled early- and late-replicating regions in Fig. 3B). However, a small number of early-firing origins do not follow the expected trend, and are present at higher copy number under limiting Pol ⍺ conditions (Fig. 3B). The three such origins on chromosome 4 are among the earliest replicating sites in the genome (Raghuraman et al., 2001), and we reasoned that the increased peak heights at these origins under limiting Pol ⍺ conditions might represent a population of cells stuck in early S-phase, corresponding to a persistent peak offset from G1 observed at 0.014% and 0.005% galactose in our flow cytometry data (Fig. S3D). To test this hypothesis, we compared DNA abundance around replication origins across our samples. While the overall abundance of genomic DNA around all replication origins was similar for 0.05%, 0.014% and 0.005% galactose (Fig. 3B), the earliest 22 origins with T_rep_ under 18 minutes (Raghuraman et al., 2001) showed increased signal at low galactose while the remaining 261 origins showed slightly decreased enrichment (Fig. 3C). Thus, for cells progressing through S-phase, our copy-number data support a global reduction in origin efficiency. Moreover, the observation that early-replicating regions are more highly sequenced than late-replicating regions across all galactose concentrations suggests that relative origin timing is unaffected by Pol ⍺ concentration.

### Ctf4 stimulates Pol α recruitment for Okazaki fragment initiation at moderate Pol ⍺ levels, and is required for efficient origin firing during severe Pol ⍺ depletion

Ctf4 has previously been shown to stimulate the recruitment or maintenance of Pol α and several additional proteins to the replisome (Gambus et al., 2009; Samora et al., 2016; Tanaka et al., 2009; Villa et al., 2016). Indeed, it has been proposed that a Ctf4 homotrimer could simultaneously recruit two Pol α complexes while the third subunit is tethered to the replisome via interaction with the Sld5 component of the Cdc45/MCM2-7/GINS (CMG) complex (Simon et al., 2014). *CTF4* is nonessential in *S. cerevisiae*; the absence of Ctf4 does not affect lagging-strand synthesis in a reconstituted *S. cerevisiae* replication system (Kurat et al., 2017), and Okazaki fragment length is unchanged in otherwise wild-type *ctf4Δ* cells (Borges et al., 2013). Pol1 contains a Ctf4 interacting peptide (CIP) motif that is necessary for Ctf4 to recruit Pol α to the fork: mutating specific residues in the CIP abolishes the interaction between Pol1 and Ctf4 without affecting the recruitment of other proteins to the fork (Simon et al., 2014, Villa et al., 2016). To investigate the effect of Ctf4 on leading- and lagging-strand synthesis under limiting Pol α conditions, we abrogated the Pol α-Ctf4 interaction via deletion of *CTF4* or mutation of the Pol1 CIP box (*pol1-4A*) (Simon et al., 2014). We analyzed Okazaki fragments from these strains using, end labeling, Southern blots, and sequencing.

In the absence of Ctf4, Okazaki fragments increase in length at ∼0.06% galactose (Fig. 4A,)–significantly higher than the 0.014% observed for a *CTF4* wild-type strain (cf. Fig. 1D&E). The *GDpol1-4A* strain showed a similar increase in Okazaki fragment length to the *ctf4Δ* strain, with longer fragments observed at 0.05% galactose (Fig. 3B&C, replicate experiments in Fig. S5A&C). These longer Okazaki fragments at sub-endogenous Pol1 levels (0.05%) were further confirmed by Southern blot with a whole genome probe (Fig. S5B). Continued growth while synthesizing long Okazaki fragments was observed down to 0.005% galactose (Fig. 4B&C). It has previously been demonstrated that overall cellular levels of Pol1 are unaffected by the absence of, or a failure to bind, Ctf4 (Villa et al., 2016). Our data therefore suggest that Ctf4 helps to maintain robust lagging-strand priming when Pol α activity is reduced, although cellular levels of Pol α are sufficient for normal Okazaki fragment initiation even in its absence (Fig. 4A and (Borges et al., 2013)).

**Figure 4.**
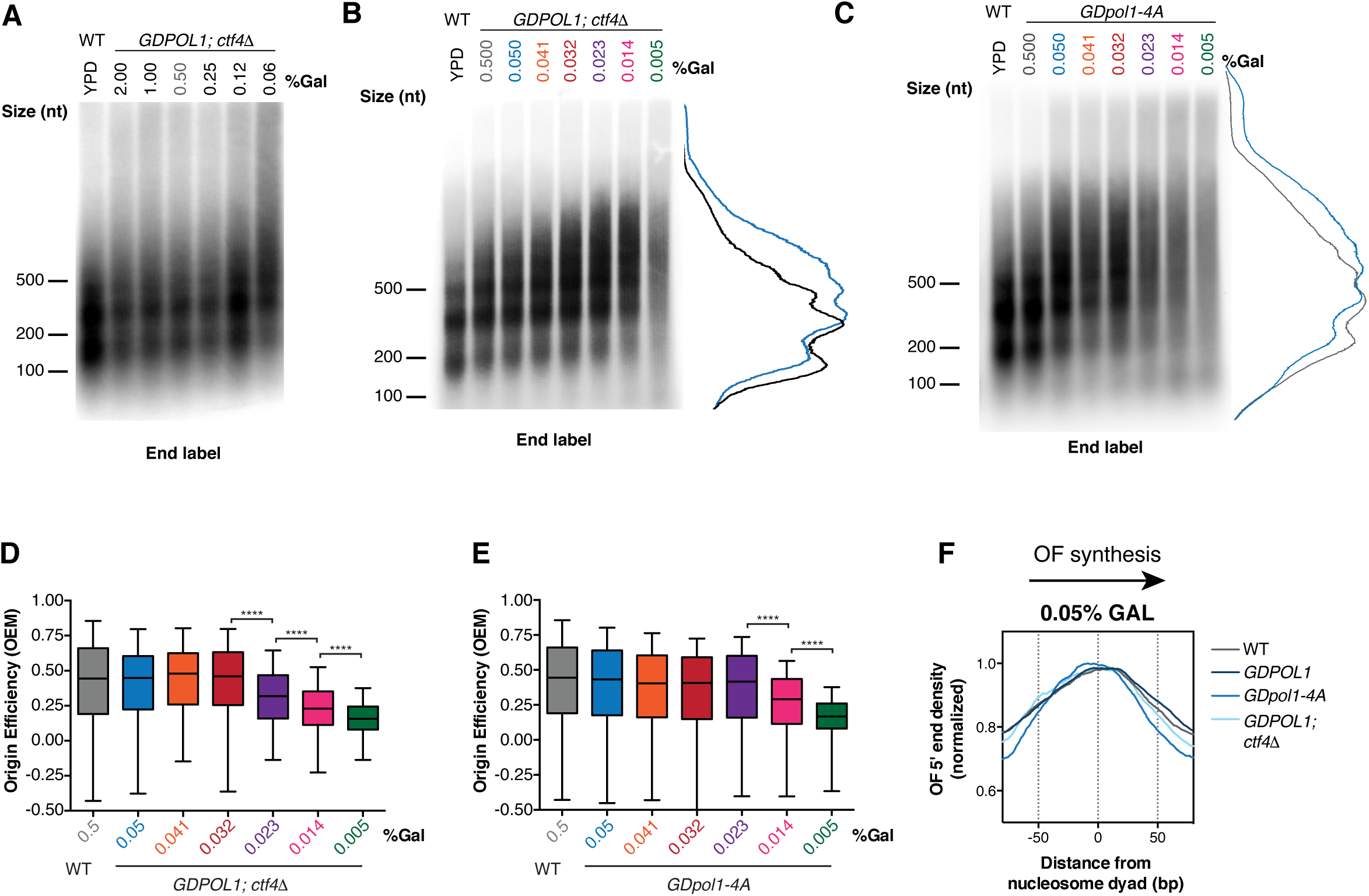
Ctf4 maintains Okazaki fragment length at moderate Pol α concentrations but is dispensable for origin firing *in vivo* unless Pol ⍺ is severely limiting. **(A, B, C).** Alkaline agarose gel analysis of end-labeled Okazaki fragments from a wild type or *GDPOL1;ctf4Δ* strain (A&B), or *GDpol1-4A* (C) as indicated, shifted to media containing various concentrations of galactose. Note that the range of galactose concentrations in A is significantly higher than in other figure panels. Traces of lanes are shown for wild type YPD (black), or *GDPOL1* 0.5% (gray) and 0.05% galactose (blue). **(D, E).** Replication origin efficiency, as in Fig. 2B, for *GDPOL1;ctf4Δ* (D) or *GDpol1-4A* (E) cells at low galactose concentrations. Data were calculated and analyzed as in Fig. 2B. **(F).** Distribution of Okazaki fragment 5’ termini around consensus nucleosome dyads, as in Fig. 1F,G, for each indicated strain shifted to media containing 0.05% galactose. Data were calculated and analyzed as in Fig. 1F&G.

We analyzed replication origin firing efficiency in the absence of Ctf4-Pol α interactions by sequencing Okazaki fragments from *ctf4Δ;GDPOL1* and *GDpol1-4A s*trains grown at various galactose concentrations (Fig. 4C). Data shown are the average of two replicates. Correlations between replicates were extremely robust at all concentrations of Pol α (Fig. S6A&B). In the absence of Ctf4, cells maintained normal levels of origin firing down to 0.032% galactose (Fig. 4D). The *GDpol1-4A* strain showed normal origin efficiency down to 0.023% galactose (Fig. 4E). Neither Fkh status (Knott et al., 2012) nor firing time (Raghuraman et al., 2001) modulates the impact of Ctf4 on origin efficiency (Fig. S7A-C). Notably, in both *ctf4Δ;GDPOL1* and *GDpol1-4A s*trains, further reduction in Pol1 levels below the initial threshold for decreased origin efficiency led to progressively lower origin firing (Fig. 4D&E). This additional decrease in origin efficiency was not observed in the *GDPOL1* strain with wild-type *CTF4* (Fig. 2B). Therefore, while the absence of Ctf4 does not appear to impact origin firing at moderate levels of Pol ⍺, the Ctf4-Pol1 interaction appears to maintain relatively robust origin firing when Pol ⍺ is severely depleted. We conclude that Ctf4-mediated recruitment of Pol α to the replisome does not stimulate replication-origin firing in *S. cerevisiae* unless Pol ⍺ is severely limiting, but plays an important role in maintaining the robustness of lagging-strand priming to fluctuations in the availability of Pol α.

Each rDNA repeat in *S. cerevisiae* contains a replication origin. rDNA size has been reported to change due to deletion of *CTF4* (Sasaki and Kobayashi, 2017) or lithium acetate transformation (Kwan et al., 2016). Expansion of the rDNA repeat increases the total number of origins in the genome, and could thereby depress origin firing elsewhere (Shyian et al., 2016; Yoshida et al., 2014). We investigated whether the origin efficiency in *ctf4Δ* cells is impacted by the size of the rDNA array. The proportion of Okazaki fragments mapping to the rDNA in *ctf4Δ* libraries was around 70% higher than in *CTF4* or *pol1-4A* libraries (Fig. S5E), consistent with substantial array expansion in the absence of Ctf4 (Sasaki and Kobayashi, 2017). However, this change in rDNA copy number appears to have a minimal effect on genome-wide origin firing efficiency.

To address the possibility that chromatin assembly defects may be the major cause of longer Okazaki fragments upon Pol α depletion, we analyzed the distribution of Okazaki fragment termini around nucleosome dyads at 0.05% galactose – a concentration that generates longer Okazaki fragments only in the context of the *ctf4Δ;GDPOL1* and *GDpol1-4A s*trains (Fig. 3A-C) but not the wild-type strain (Fig. 1C). At this intermediate Pol α concentration, the distribution of Okazaki fragment 5’ and 3’ termini was highly nucleosome-biased and very similar between wild-type, *GDPOL1, ctf4Δ;GDPOL1,* and *GDpol1-4A s*trains (Fig. 3F & S5F-G). At low Pol ⍺ concentrations, the distribution of Okazaki fragment 5’ and 3’ termini lost nucleosome patterning in *ctf4Δ;GDPOL1,* and *GDpol1-4A* (Fig. S5F-G), similarly to the behavior of the wild-type *GDPOL1* strain (Fig. 1G). However, the alignment of Okazaki fragment 5’ or 3’ end locations with nucleosome dyads was lost at higher Pol ⍺ concentrations in *ctf4Δ;GDPOL1* strains compared to *GDpol1-4A* (Fig. S5F-G), consistent with an additional contribution of Ctf4 to chromatin assembly beyond Pol ⍺ recruitment.

*ctf4Δ;GDPOL1,* and *GDpol1-4A* cells grown at 0.05% galactose show increased Okazaki fragment length (Fig. 4B-C) but no defect in the nucleosome patterning of Okazaki fragment termini (Fig. 4F). Therefore, Okazaki fragment length can be increased by reduced Okazaki fragment initiation in the absence of an accompanying chromatin assembly defect. We note that these data are not inconsistent with impaired nucleosome assembly at severely reduced Pol α concentrations contributing to a further increase in Okazaki fragment length.

### Checkpoint activation is required for viability when origin firing is reduced, but not when lagging-strand priming is perturbed

In response to DNA damage or replication stress, a checkpoint signaling cascade is initiated and culminates in the phosphorylation of the effector kinase Rad53 (Ciccia and Elledge, 2010). We analyzed Rad53 phosphorylation in the *GDPOL1* strain after a switch to low galactose. Substantial phosphorylation of Rad53 was observed only at concentrations below 0.023% galactose (Fig. 5A) – a concentration at which both origin firing and Okazaki fragment initiation are perturbed (Figs. 1&2). To further investigate the interplay between Pol α depletion, increased Okazaki fragment length, decreased origin firing, and checkpoint activation, we combined the *GDPOL1* allele with deletion of *MEC1* (with additional deletion of *SML1* to maintain viability of *mec1Δ* cells). Growth of the *GDPOL1; mec1Δ; sml1Δ* mutant was impaired at 0.05% galactose and virtually absent at 0.014% and 0.005% galactose (Fig. 5B). Thus, checkpoint-deficient cells cannot survive with limiting Pol α. Consistent with a rapid loss of viability upon Pol1 depletion in checkpoint-deficient cells, we were unable to robustly detect long Okazaki fragments in *GDPOL1; mec1Δ; sml1Δ* cells shifted to low concentrations of galactose (Fig. 5C). Okazaki fragment length was normal in this strain at galactose concentrations above 0.023%.

**Figure 5.**
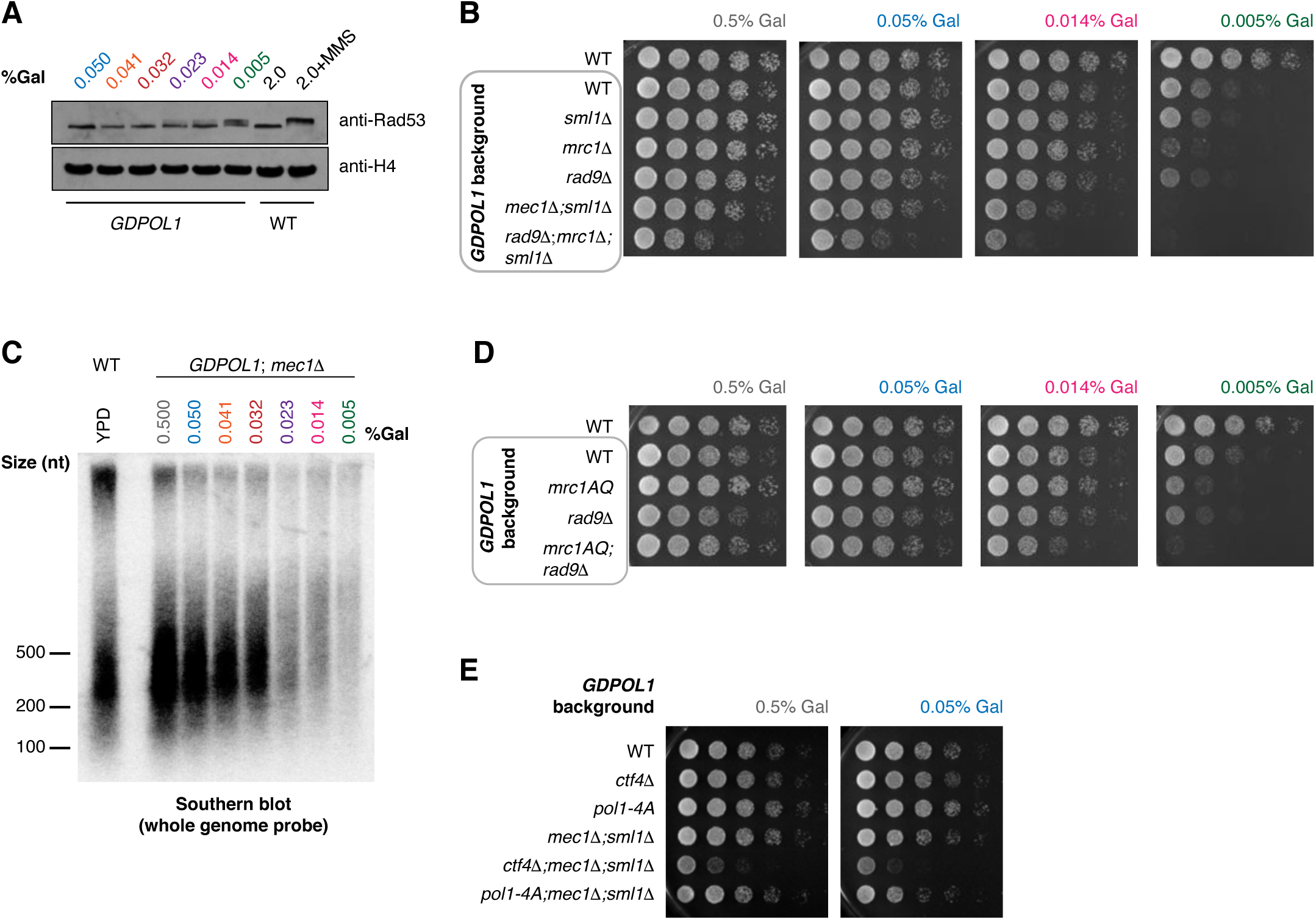
The checkpoint is required for viability under conditions of limiting origin firing, but not increased Okazaki fragment size. **(A).** Western blot against Rad53 from asynchronous *GDPOL1* cells shifted to the indicated media. A wild-type strain grown in YPGal ± 0.1% MMS were used as negative and positive controls for Rad53 hyperphosphorylation. **(B).** Serial dilution spot tests to assay the growth of *GDPOL1* strains carrying additional mutations (*mec1Δ;sml1Δ, rad9Δ, mrc1Δ, rad9Δ;mrc1Δ;sml1Δ*) at the indicated galactose concentrations. **(C).** Southern blot as in Fig. 3B, for a *mec1Δ;GDPOL1* strain shifted to various galactose concentrations. **(D).** Serial dilution spot tests to assay the growth of *GDPOL1* strains carrying additional mutations (*rad9Δ, mrc1^AQ,^ rad9Δ;mrc1^AQ^*) at the indicated galactose concentrations. **(E).** Serial dilution spot tests to assay the growth of *ctf4Δ;GDPOL1* or *GDpol1-4A* strains with or without additional deletion of *MEC1*.

To test the contributions of the DNA damage checkpoint (DDC), mediated by Rad9, and the DNA replication checkpoint (DRC), mediated by Mrc1, to survival under limiting Pol α conditions, we analyzed the growth of *GDPOL1* strains in combination with *rad9Δ*, *mrc1Δ*, or the non-phosphorylatable *mrc1^AQ^* allele (Osborn and Elledge, 2003) (Fig. 5B&D; the full range of galactose concentrations is shown in Fig. S8A-B). *GDPOL1* cells showed robust growth at low Pol1 concentrations (0.014% galactose) in both *rad9Δ* and *mrc1Δ* strain backgrounds. *GDPOL1;mrc1^AQ^; rad9Δ* cells did not grow at 0.005% galactose (Fig. 5D), confirming that the inviability of *GDPOL1; mec1Δ; sml1Δ* cells under these conditions is due to the absence of functional DDC and DRC signaling. We note that *GDPOL1;mrc1^AQ^* cells grow better than *GDPOL1;mrc1Δ* cells at all concentrations tested; this suggests that the slight growth defect at 0.005% galactose observed in *GDPOL1;mrc1Δ* cells is most likely due to decreased replication fork speed (Szyjka et al., 2005; Yeeles et al., 2017) in combination with decreased origin firing (Fig. 2B) as opposed to impaired DRC activity. In summary, checkpoint activation is required for viability under conditions of severe Pol α depletion: this activation can proceed via either the DDC or the DRC.

We investigated the sensitivity of *GDPOL1* cells during Pol α depletion to either replication stress or DNA damage by testing growth defects in the presence of 1 mM hydroxyurea or 0.008% methyl methanesulfonate (Fig. S8A-D). We observed that growth defects in the presence of HU or MMS were exacerbated by lowering the concentration of Pol ⍺; this effect was apparent in both checkpoint-proficient and checkpoint-deficient strains. Indeed, in the presence of 1 mM HU a significant growth defect can be observed at 0.05% galactose – a concentration at which neither origin firing nor lagging-strand priming is impaired in strains proficient for Ctf4-Pol ⍺ interaction.

To determine whether the requirement for checkpoint activation was due to deregulated lagging-strand priming or impaired leading-strand initiation, we compared the growth of *GDPOL1;mec1Δ;sml1Δ* strains with and without *CTF4* or *pol1-4A* at 0.5% and 0.05% galactose. Loss of Ctf4-Pol α interactions affects Okazaki fragment initiation at relatively high Pol α concentrations, before leading-strand initiation is impaired. Thus, *ctf4Δ* and *pol1-4A* are effectively separation of function mutants: at 0.05% galactose, Okazaki fragment initiation is reduced by the absence of Ctf4-mediated recruitment of Pol α while leading-strand initiation is not (Fig. 4). *GDPOL1;mec1Δ;sml1Δ;ctf4Δ* cells grow slowly relative to *MEC1* or *CTF4* cells: however, growth was minimally affected by galactose concentrations (Fig. 5E). This result is recapitulated with *GDpol1-4A;mec1Δ;sml1Δ* strain, which showed essentially no growth defect at either Pol α concentration (Fig. 5E). Therefore, perturbed lagging-strand synthesis does not directly cause growth defects the absence of a functional checkpoint. We conclude that cells with limiting Pol α become reliant on the checkpoint at least in part due to decreased replication origin firing as opposed to solely due to increased Okazaki fragment length.

## Discussion

### Cellular impact of perturbed leading- and lagging-strand initiation

Checkpoint activation is required for robust DNA synthesis, and therefore for viability, when origin firing is significantly reduced (Fig. 5). This observation is consistent with genetic interactions in *S. cerevisiae* – for example similar negative genetic interactions of *rad9Δ* with alleles of the catalytic subunits of each of the three replicative polymerases (Dubarry et al., 2015). It will be interesting to determine the contributions of leading- and lagging-strand perturbations to the many reported phenotypes resulting from *pol1* mutation or Pol1 depletion – for example increased trinucleotide repeat expansion rate and size (Shah et al., 2012), and increased chromosome fragility (Lemoine et al., 2005; Song et al., 2014). Furthermore, mutations that reduce the levels of functional Pol α (Van Esch et al., 2019) or Pol ε (Bellelli et al., 2018) in mammalian cells increase replication stress linked to reduced origin firing, highlighting the relevance of these studies beyond budding yeast.

### Pol α initiation on the leading and lagging strands

Our data indicate that both origin firing and Okazaki fragment initiation in *S. cerevisiae* are robust with respect to fluctuations in Pol α availability. Pol α concentration does not normally limit primer synthesis and/or utilization, and significant disruption of Okazaki fragment synthesis or replication initiation is only observed at very low Pol α concentrations (Fig. 1). Because cells with limiting Pol α synthesize longer Okazaki fragments, as opposed to fewer Okazaki fragments of normal size, our data suggest that Pol α can function distributively *in vivo* as opposed to being obligately tethered at the replication fork. We additionally note that the increase in Okazaki fragment length upon Pol α depletion is consistent with a priming mechanism in which there is no strict coupling between DNA unwinding and primer synthesis *in vivo* (Fig. 6A).

**Figure 6.**
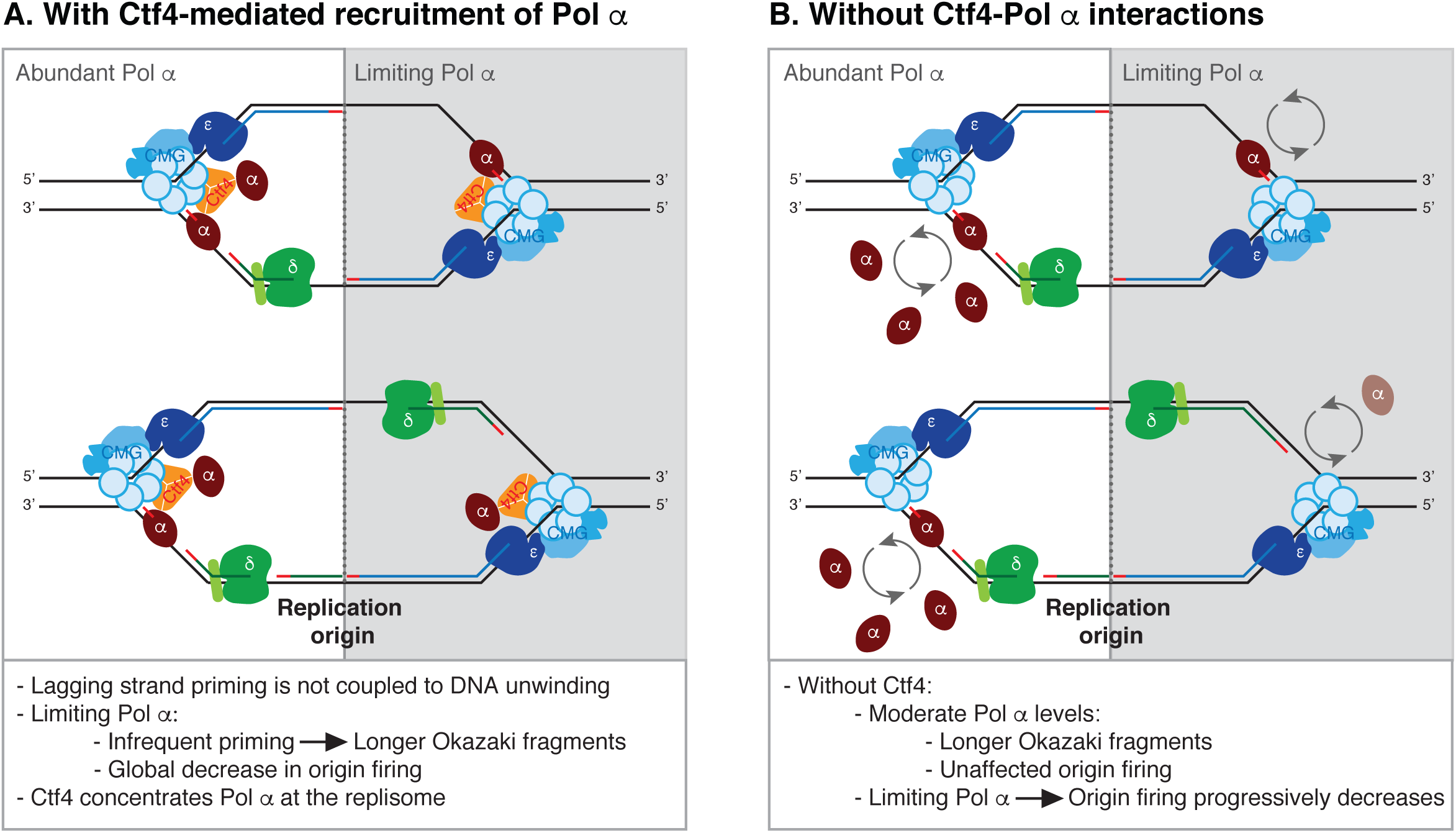
Model of Pol α recruitment for leading- and lagging-strand priming in *S. cerevisiae*

Although the Ctf4 protein has little to no effect on lagging-strand synthesis in chromatinized reconstituted systems (Kurat et al., 2017), we observe that Ctf4 stimulates lagging-strand priming *in vivo*. In the absence of Ctf4, Okazaki fragment length can still be modulated by Pol α concentration (Fig. 4). Thus, in light of the potentially distributive action of Pol α, we propose a simple model that Ctf4 acts by increasing the local concentration of Pol α at the elongating replication fork. An alternative possibility is that Ctf4 maintains two copies of Pol α at each replisome under normal conditions (Fig. 6B). It is unclear what underlies the differential Ctf4-sensitivity of reconstituted and *in vivo* lagging-strand synthesis. Our data also show that, unlike ongoing Okazaki fragment priming during lagging-strand synthesis, the productive initiation of DNA synthesis at replication origins is not impacted by the absence of Ctf4-mediated recruitment until severely limiting Pol α concentrations (Figs. 2&4). A recent report demonstrated that leading-strand initiation can occur via extension of the first Okazaki fragment from the opposite replication fork, and that the two replisomes are inter-dependent during this establishment phase (Aria and Yeeles, 2018). The close proximity of the two replisomes at this stage could underlie the differential Ctf4-sensitivity of replication-origin firing at moderate and low Pol α concentrations.

All replication origins are affected by Pol α depletion, without dependence on their normal firing time or efficiency (Fig. 2E & 3B). Therefore, it is unlikely that the reduction in origin firing is a direct result of checkpoint activation. Our data suggest that the origin firing program is determined by the relative accessibility of licensed origins to limiting soluble firing factors, even when overall origin firing is reduced. We note that coordinately down-regulating the efficiency of all origins, as opposed to selectively reducing the efficiency of a subset, represents a robust strategy to maintain the evolutionarily selected co-orientation of deleterious transcription events with replication (Chen et al., 2019; Hamperl et al., 2017; Osmundson et al., 2017; Tran et al., 2017).

### Chromatin and lagging-strand synthesis

Pol α acts distributively on naked DNA in reconstituted replication reactions (Yeeles et al., 2017). Chromatin reduces Okazaki fragment size *in vitro,* and has therefore been proposed to make Pol α more processive (Kurat et al., 2017). Our data suggest that Pol α can act distributively on chromatin *in vivo*, but is present at saturating concentrations that would minimize the difference between distributive or processive activity. Our data do, however, support an intimate interaction between chromatin and lagging-strand priming. Extreme depletion of Pol α leads to Okazaki fragment distributions consistent with impaired chromatin assembly (Fig. 1G and S5F-G). Since Ctf4 connects Pol α to the replicative helicase, its absence also impacts chromatin dynamics during DNA replication (Evrin et al., 2018). However, perturbed Okazaki fragment initiation in the absence of Ctf4-mediated recruitment of Pol α to the fork can generate longer Okazaki fragments in the absence of an obvious chromatin assembly defect (Fig. 4F). It is also possible that the increased length of Okazaki fragments reported in histone chaperone mutants (Smith and Whitehouse, 2012; Yadav and Whitehouse, 2016) represents an underlying defect in priming.

## MATERIALS AND METHODS

### Yeast strains

All yeast strains were W303 *RAD5+*, and contained additional mutations required for anchor-away depletion of Cdc9. The genotype of the wild-type strain is *mata, tor1-1::HIS3, fpr1::NatMX4, RPL13A-2xFKBP12::TRP1, CDC9-FRB::HygMX*. The *pol1-4A* mutant from (Simon et al., 2014) was generated using the CRISPR/Cas9 system in *S. cerevisiae* as previously described (Dicarlo et al., 2013). Briefly, a guide RNA was synthesized specific to the CIP box in *POL1* that could not be cleaved if repaired with donor sequence with D141A, D142A, L144A and F147A mutations and 100 bp of homology on both sides. This created a markerless strain that was confirmed through Sanger sequencing. After transformations, strains were grown on YPD to lose the Cas9 and gRNA plasmids. Additional gene deletions, and replacement of the *POL1* promoter, were carried out by PCR-mediated replacement in a wild-type strain, and introduced into the desired background by cross.

### Cell growth, cell-cycle synchronization, and spot tests

All strains were grown at 30°C, starting in YEP with 0.5% galactose plus 3% raffinose, unless otherwise noted. To deplete Pol1 levels, cultures were sugar switched by growing overnight to log phase then washing the cells with sterile deionized water then sterile YEP before being inoculated into fresh YEP media with various galactose concentrations supplemented with 3% raffinose.

For short-term experiments, strains were grown for four hours after the sugar switch before adding rapamycin for one hour of ligase repression. Cells were collected four hours after the sugar switch for western blot analysis.

For cell-cycle synchronization, cultures were sugar-switched at log phase and then added 5g/mL alpha factor to synchronize cells in G1 phase. Cells were released into S phase and were collected every 15 minutes by centrifugation at 4°C then stored at −80°C or by immediately fixing cells in 70% ethanol.

For spot tests, yeast cells were washed then counted. Similar numbers of cells were plated onto various galactose concentrations with or without 1mM hydroxyurea (Sigma H8627) or 0.008% methyl methanesulfonate (Sigma 129925) at a 1:5 dilution series and grown overnight for two days at 30°C unless indicated otherwise.

### Fluorescence-activated cell sorter (FACS) analysis

Cells were collected after release from G1 arrest every 15 minutes and fixed in 70% ethanol and incubated at 4°C overnight. Fixed cells were then spun down and resuspended in 50mM sodium citrate with RNase A (Fisher 50-153-8126) for 1 hour at 50°C. Next, with the addition of proteinase K (MP Biomedicals) the samples were incubated for 1 hour at 50°C. Cells were then stained with SYTOX green (Fisher S7020) then sonicated and processed using a Becton Dickinson Accuri.

### Western blotting

Samples were collected by centrifugation and washed with deionized water and stored at −80°C before lysate preparation. Lysates were prepared by five-minute resuspension in 600uL 2M lithium acetate on ice, pelleted, resuspended in 600uL 400mM sodium hydroxide at room temperature for five minutes, pelleted, and resuspended in Laemmli buffer with 5% beta-mercaptoethanol prior to boiling, then briefly pelleted immediately prior to loading the lysate onto a SDS-PAGE gel. Samples were transferred to PVDF, blocked with 5% milk and probed with C-Myc antibody (Genscript A00173-100), Rad53 antibody (Abcam ab104232), histone H4 antibody (Abcam ab10158), or actin antibody (Thermofisher Scientific MA1-744).

### Okazaki fragment analysis by gel electrophoresis

Following the sugar switch methods above, Okazaki fragments were accumulated by adding rapamycin (Spectrum 41810000-2) to 1ug/mL to anchor away Cdc9, which is tagging with FRB, for 1h (Haruki et al., 2008). These cells were collected through centrifugation and immediately processed or stored at −80°C. Genomic preps from spheroplasts were completed as previously described (Smith and Whitehouse, 2012). For end-labeling, 5uL of genomic DNA was labeled using a 50uL reaction with 5U Klenow exo-(NEB M0212L) and α-dCTP (Perkin Elmer BLU513H500UC) at a final concentration of 33 nM. Reactions were then ethanol precipitated to remove excess label. Normalized amounts (from either native gel or previous experiment) were loaded onto a 1.3% denaturing agarose gel. After electrophoresis, gels were then blotted onto nitrocellulose membrane (Fisher 45-000-932) overnight. Membranes were then dried and exposed to phosphor screens.

For southern blot analysis, unlabeled genomic DNA was normalized to total genomic DNA, and run on 1.3% denaturing agarose gels. After electrophoresis, the gel was blotted onto a nitrocellulose membrane (Fisher 45-000-932) overnight. Next the membrane was crosslinked and washed, then hybridized overnight with a probe synthesized with random hexamers labeling kit (Fisher 18187-013) and sheared genomic DNA. After two low stringency washes, membranes were dried then exposed to phosphor screens.

### Okazaki fragment purification, sequencing, and analysis

Genomic DNA was boiled at 95°C for 5 min then salt was added to 300mM NaCl, pH 12. Purification of Okazaki fragments was accomplished by running the denatured genomic DNA through 400 ul Source 15Q (VWR 89128-854), binding at 300 mM NaCl, pH 12. DNA was eluted in 50mM steps until 900mM NaCl, pH 12, fractions kept were 800mM, 850mM, and 900mM, these were stored at −20°C. DNA was ethanol precipitated then treated with RNase cocktail (Thermo Fisher AM2286) for 1h at 37°C to remove any RNA. Next, these reactions were ethanol precipitated then run through Illustra microspin G-50 columns (Fisher 27-5330-02). The single-strand Okazaki fragments were boiled at 95°C for 5 min and cooled quickly on ice and up to 1ug of purified fragments were ligated with T4 DNA ligase (Fisher 50305904) to 1ug of adaptors with single-stranded overhangs that were generated as previously described (Smith and Whitehouse, 2012). Purified libraries were amplified (12-16 cycles) using Illumina Truseq primers according to Illumina protocols, but with Phusion (NEB M0530L). Paired-end sequencing (2 × 75 bp) was carried out on an Illumina Next-seq 500 platform. FASTQ files were aligned to the s288c reference genome using the Bowtie (v2.3.2). The files were converted, then bad quality reads and PCR duplicates were removed using the Samtools suite (v1.9). Then the genomic coverage was calculated using the Bedtools suite (v2.27.1) in a strand-specific manner to make stranded bed files. Origin efficiency metric analysis was achieved through calculating the strand bias in 10kb windows around predefined origins as previously described (McGuffee et al., 2013) with the origin list from the same source. To map the Okazaki fragments ends relative to known nucleosome dyads (Jiang and Pugh, 2009), 5’ and 3’ fragment ends were extracted from previously generated bed files and a meta-analysis was completed as previously described (Smith and Whitehouse, 2012).

### Whole genome sequencing and analysis

Cells were incubated for 4h at various galactose concentrations with a simultaneous arrest with 5mg/ml alpha factor for 3h. Cells were collected every ten minutes after a room temperature (25°C) release. Samples were collected and stored for flow cytometry at 4°C and for whole genome sequencing at −80°C. Samples were selected for analysis based on flow cytometry data. All samples, including a G1 control, were lysed using a FastPrep system. The lysates were then sonicated using a Branson 250 sonicator at 15% for 15 seconds 5X. The sheared DNA was then treated with 100μg of Proteinase K for 2h at 37°C. Samples were then phenol chloroform extracted and precipitated. DNA was then quantified and libraries were prepped using TruSeq Nano DNA LT Kit (Illumina 20015964). Sequencing and alignment methods were performed as mentioned above, but sequencing was not strand-specific. Pertinent S phase samples were pooled for analysis. These genomic coverage files were normalized to the median coverage, binned to 100bps, and poorly mapped sites were removed from all datasets (Table S2). Next, the coverage was calculated by taking the read depth normalized to G1 and the smoothed using a Loess regression in R. To perform a meta-analysis around all or specific origins of replication, the average coverage around these origins were calculated using the same origin list as above.

## ACKNOWLEDGEMENTS

We thank the NYU Gencore for assistance with TapeStation and sequencing, and members of the Smith lab for helpful discussions.

## LEGENDS TO SUPPLEMENTARY FIGURES

**Figure S1.**
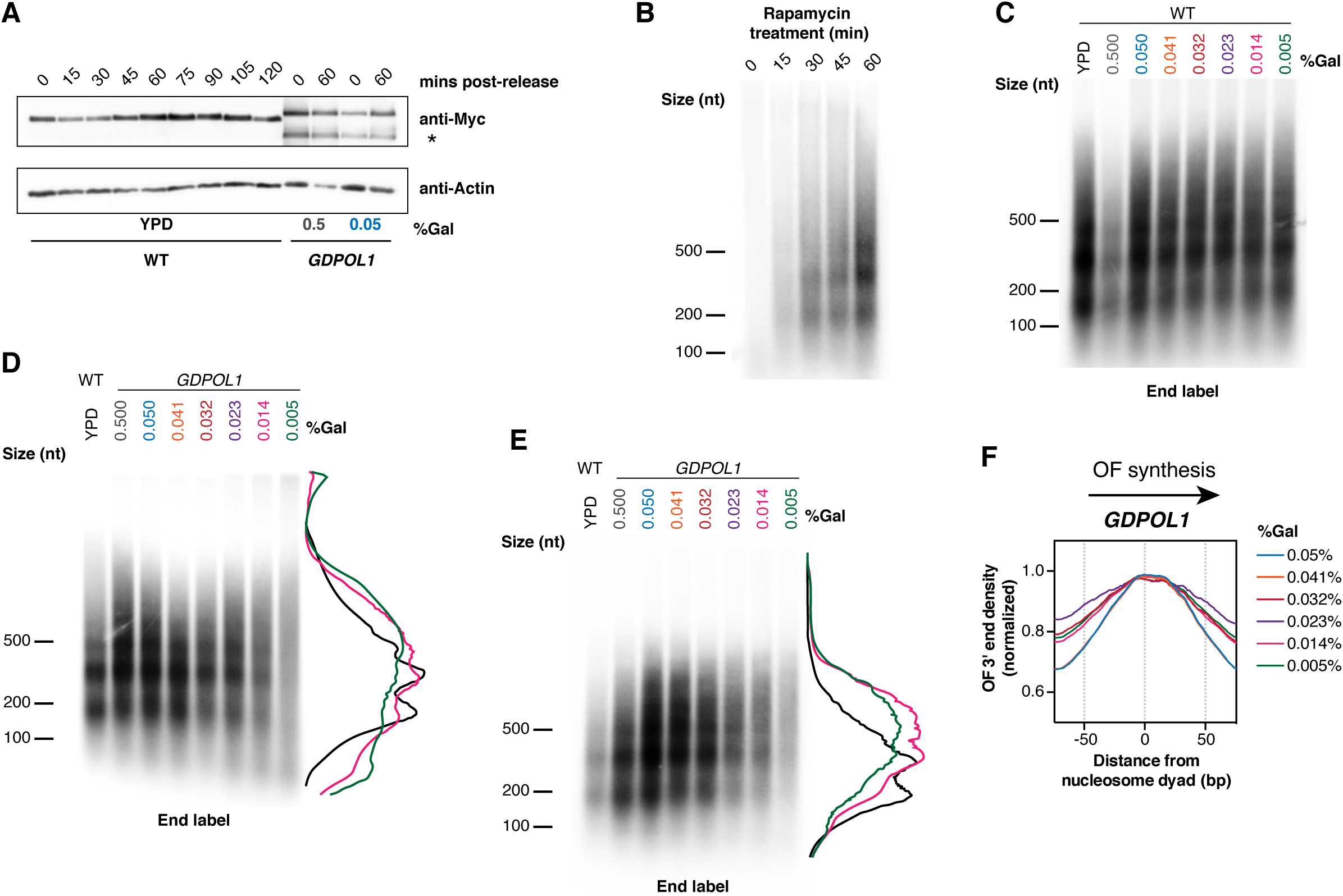
(Associated with Fig 1) **(A).** Western blot against 13xMyc-tagged Pol1 from wild-type or *GDPOL1* cells at the indicated sugar concentration, released from alpha-factor arrest for the indicated time. The GDPol1-specific degradation product is indicated by an asterisk. **(B).** Timecourse of Okazaki fragment enrichment during Cdc9 nuclear depletion by anchor away (Haruki et al., 2008). Okazaki fragments were prepared and labeled as in Fig. 1C, and as previously described (Smith and Whitehouse, 2012). **(C-E).** Representative replicate Okazaki fragment end-labeling gels for wild-type (C) and *GDPOL1* (D-E) at the indicated galactose concentrations. Traces adjacent to the plots in D&E indicate the change in size distribution of Okazaki fragments at 0.014% (pink) and 0.005% (green) galactose. **(F).** Distribution of Okazaki fragment 3’ ends around consensus nucleosome dyads (Jiang and Pugh, 2009) in the *GDPOL1* strain shifted to media containing the indicated concentration of galactose

**Figure S2.**
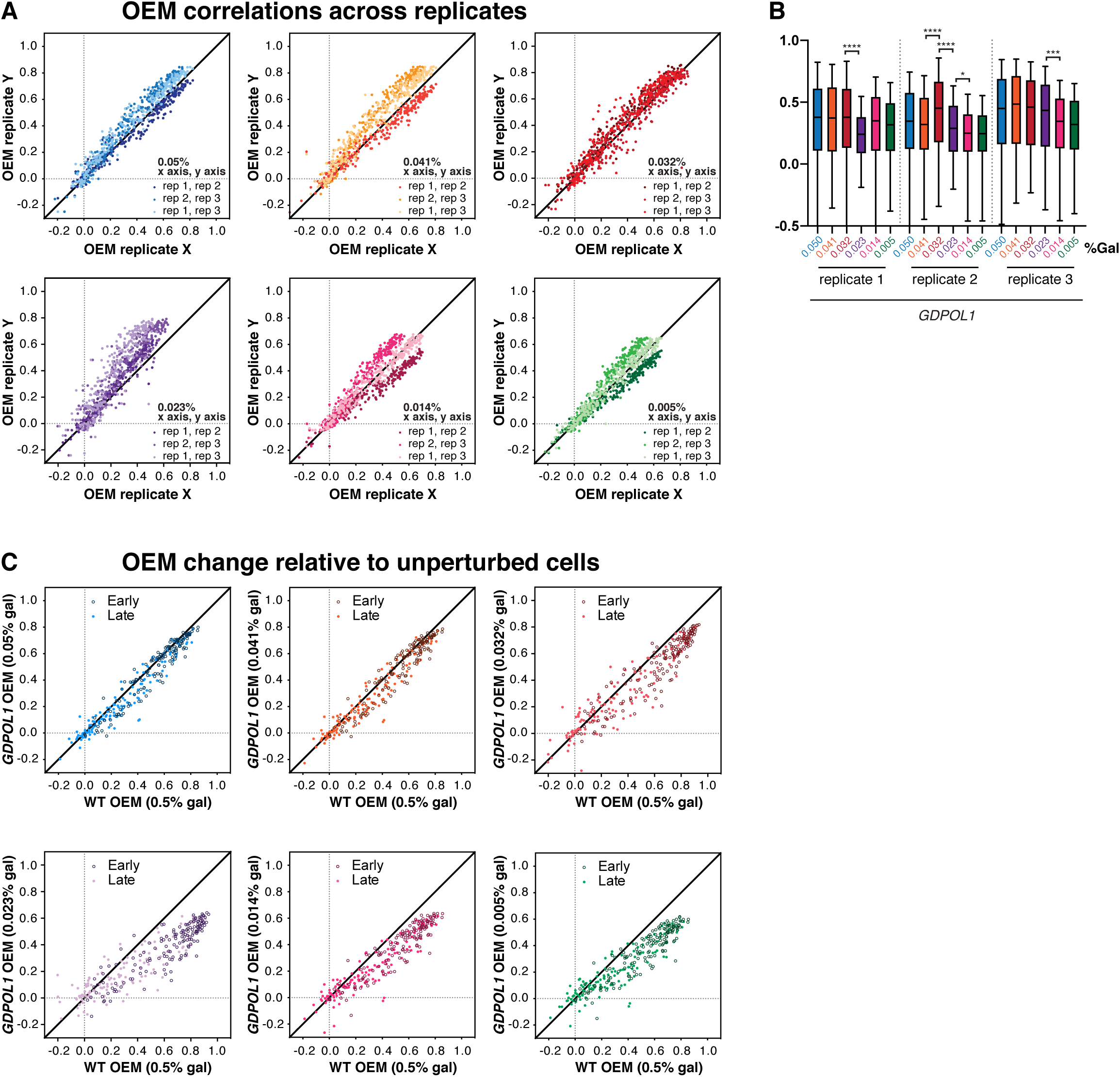
(Associated with Fig 2) **(A).** Origin efficiency replicate comparisons for data from the *GDPOL1* strain shown in Fig. 2B each of three replicates are plotted against each other and indicated by color. **(B).** Comparison of origin efficiency data from each replicate from *GDPOL1* cells shifted to the indicated concentration of galactose. Significance was calculated by unpaired t-test; **** p<0.0001, * p<0.05. **(C).** Scatter plots comparing origin firing efficiency in *GDPOL1* cells at various galactose concentrations, to wild-type cells in 0.5% galactose.

**Figure S3.**
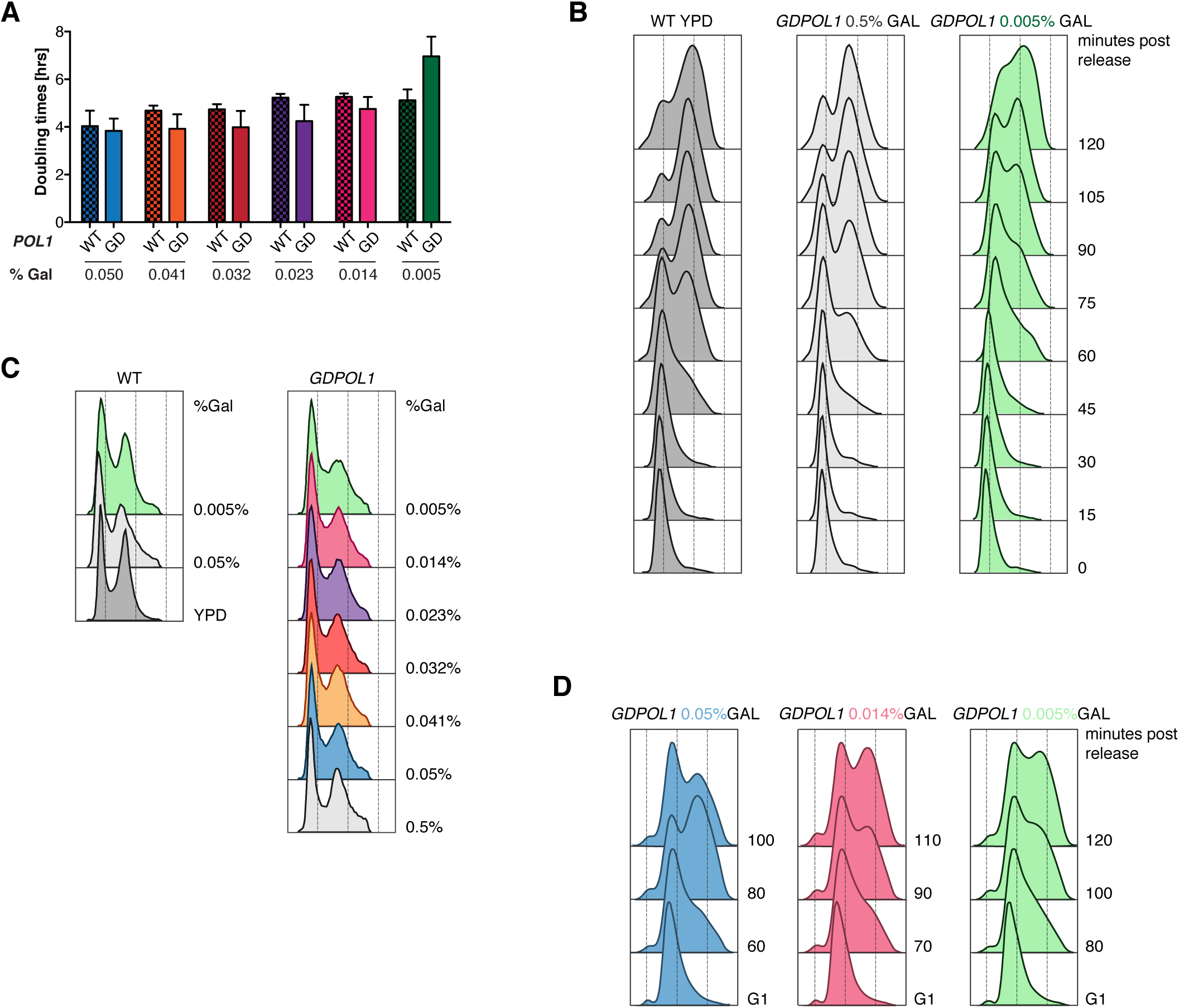
(Associated with Fig 2) **(A).** Doubling times for wild-type or *GDPOL1* strains in YEP + 3% raffinose, supplemented with the indicated concentration of galactose. Data are the average of at least three replicates in each case. **(B).** DNA content, assayed by flow cytometry, of an arrest release of wild type or *GDPOL1* cells also analyzed in A. **(C).** DNA content, assayed by flow cytometry, of asynchronous cells post 4h sugar switch of wild type or *GDPOL1* cells. **(D).** DNA content, assayed by flow cytometry, *GDPOL1* cells, released into S-phase after 4h sugar switch in G1. The samples collected at these time points were used to generate sequencing libraries for the analysis shown in Fig. 3.

**Figure S4.**
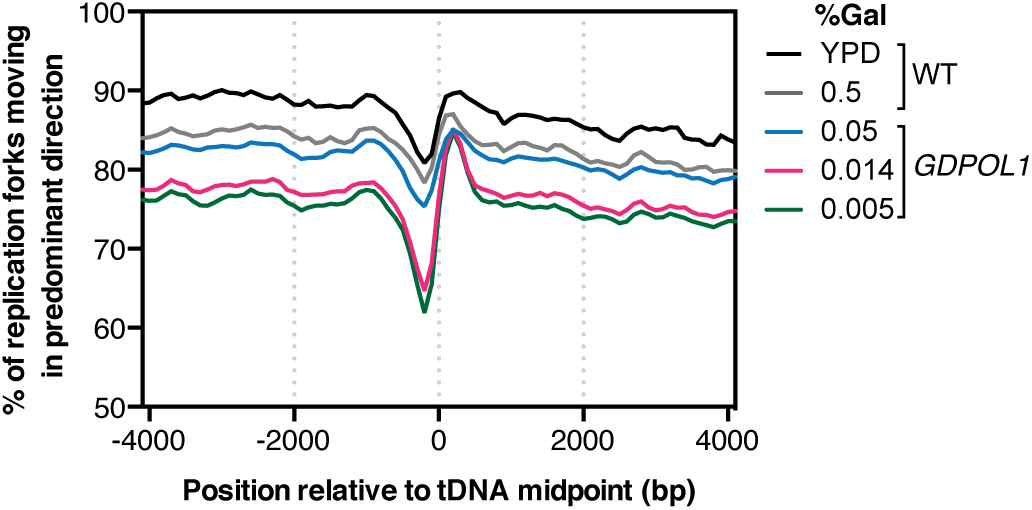
Analysis of replication-fork direction around 93 origin-distal tRNA genes (Osmundson et al., 2017) in *GDPOL1* cells grown at various galactose concentrations. Increased replication-fork stalling or arrest at these sites would manifest as a decrease at or after the midpoint of the gene (Osmundson et al., 2017).

**Figure S5.**
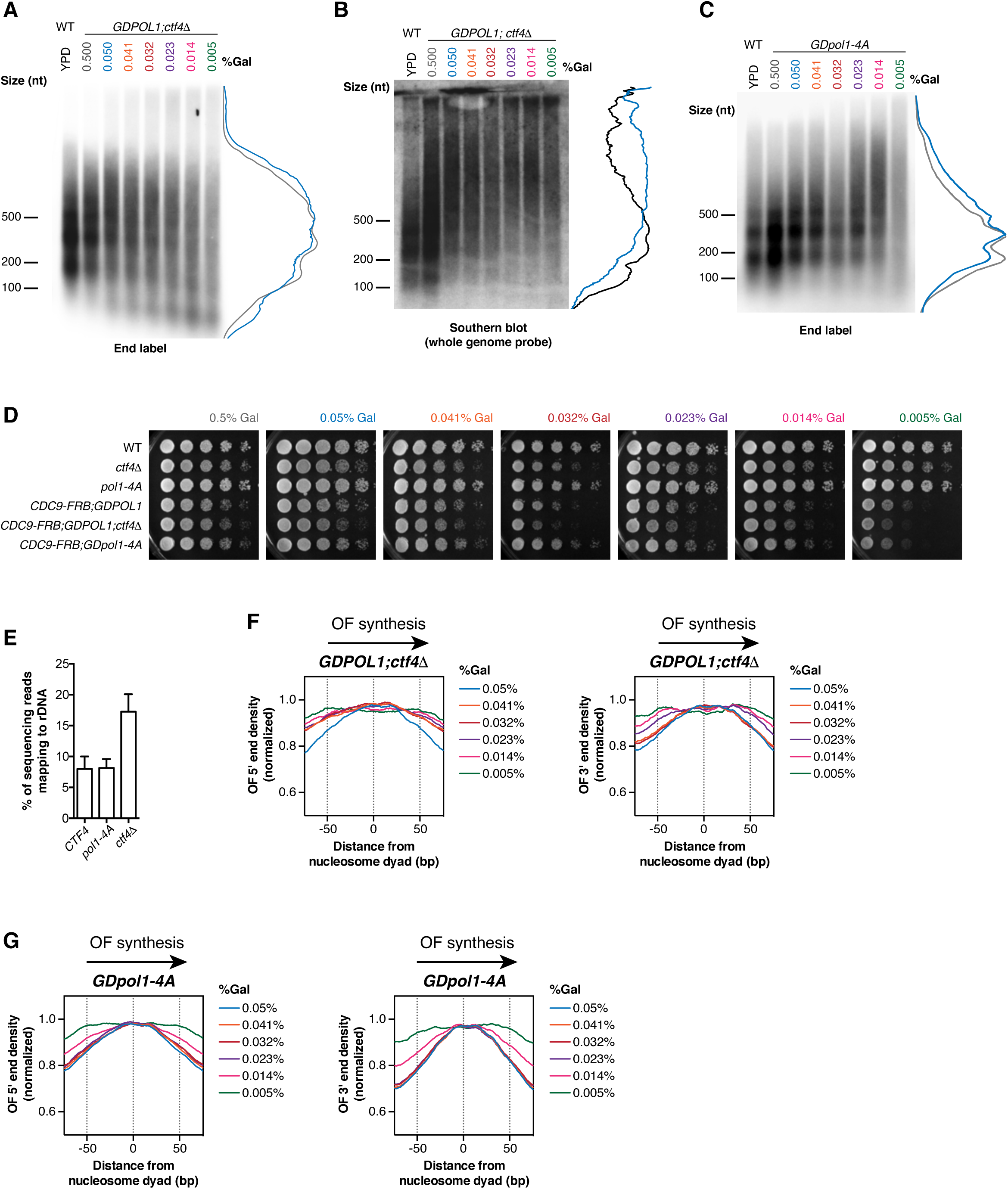
(Associated with Fig 3) **(A-C).** Representative replicate end-labeling gel (A, C) or Southern blot (B), on Okazaki fragments from a *GDPOL1;ctf4Δ* (A, B) or *GDpol1-4A* (C) strain shifted to media shifted to low galactose concentrations. Traces of YPD (black) and 0.05% galactose (blue) lanes on the right. A control lane for wild-type cells grown in YPD is included on each gel. **(D).** Serial dilution spot tests to assay the growth of *GDPOL1* strains with or without FRB tagging of CDC9 and/or *ctf4Δ* or *pol1-4A* mutations. **(E).** The rDNA repeat is expanded in *ctf4Δ; GDPOL1* cells. The proportion of sequencing reads mapping to the rDNA is indicated. Data represent the mean ± SD of all sequencing datasets used for analysis in figures 2&3. **(F-G).** Distribution of Okazaki fragment 5’ (left panel) and 3’ ends (right panel) around consensus nucleosome dyads (Jiang and Pugh, 2009) in the *GDPOL1;ctf4Δ* (F) or *GDpol1-4A* strain (G) shifted to media containing various galactose concentrations.

**Figure S6.**
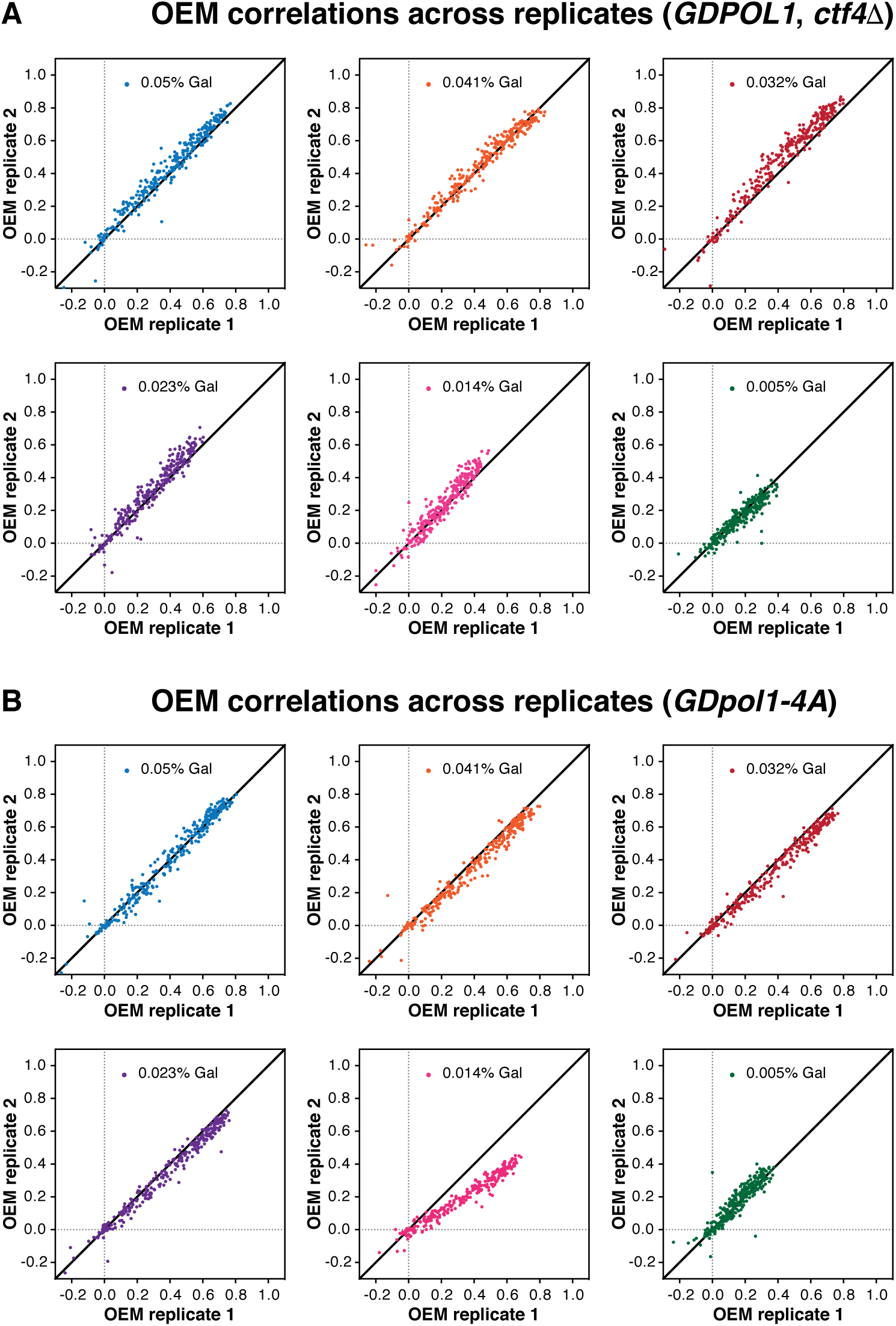
(Associated with Fig 3) **(A, B).** Origin efficiency replicate comparisons for data from the *ctf4Δ;GDPOL1* strain shown in Fig. 3D (A) and the *GDpol1-4A* strain show in Fig. 3F (B).

**Figure S7.**
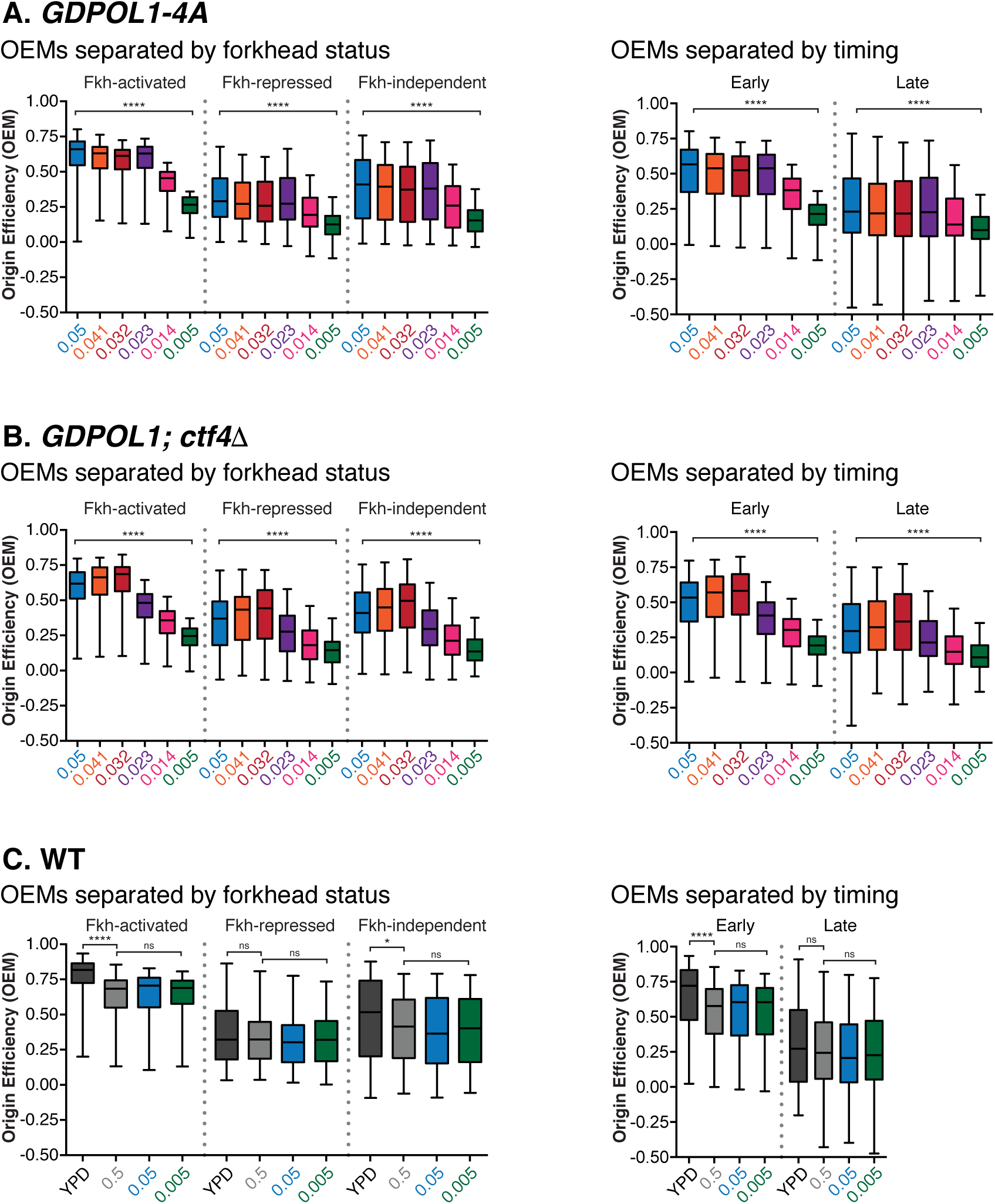
(Associated with Fig 4) **(A, B, C).** Firing efficiency for origins separated by Fkh status or replication timing for the data sets in Fig. 4. Significance was calculated by unpaired t-test; **** p<0.0001, * p<0.05.

**Figure S8.**
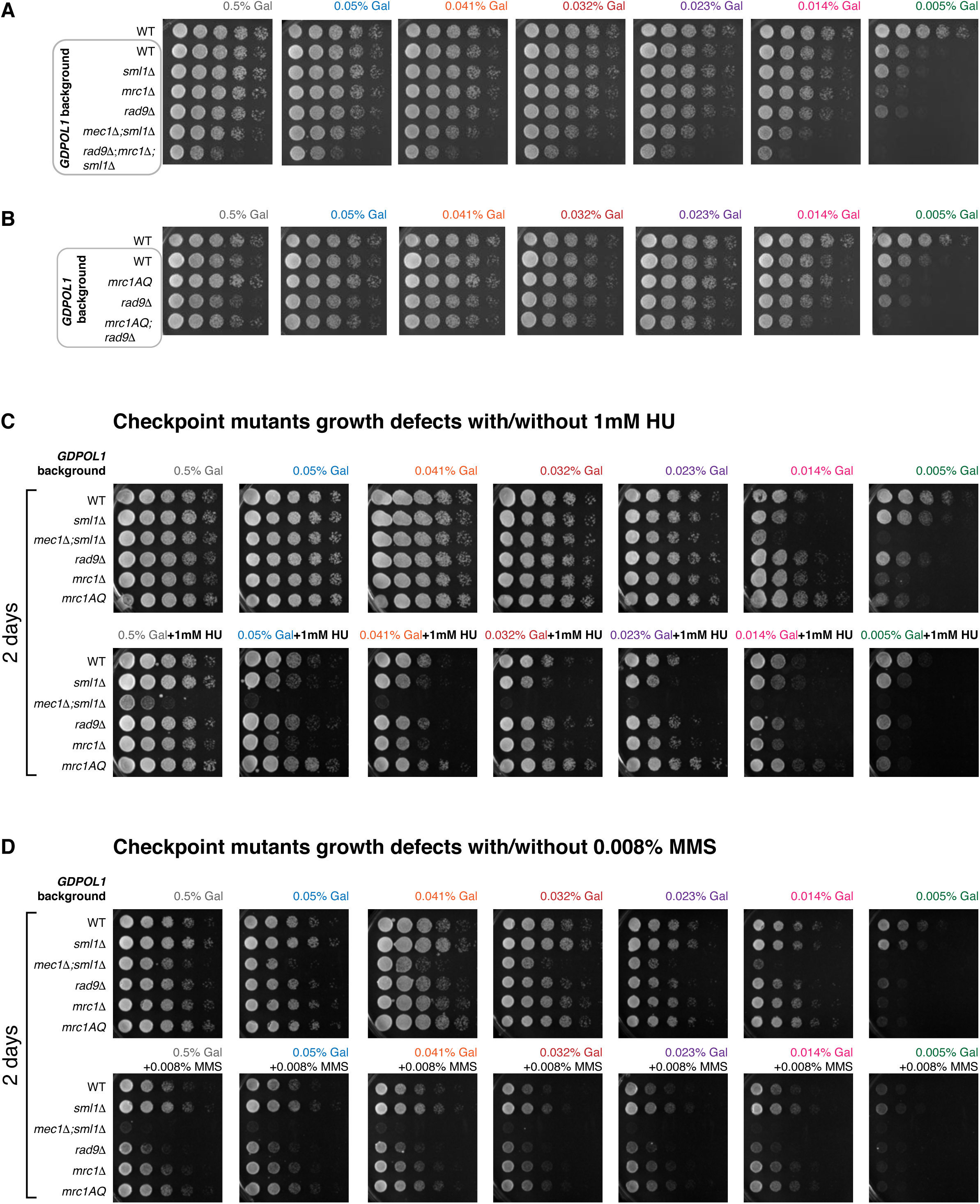
(Associated with Fig 4) **(A, B).** Serial dilution spot tests to assay the growth of *GDPOL1* strains carrying additional mutations (*mec1Δ;sml1Δ, rad9Δ, mrc1Δ, mrc1AQ*) at the indicated galactose concentrations. A selection of these concentrations is shown in Fig. 4A-B **(C,D).** Serial dilution spot tests of the indicated strains with or without 1 mM hydroxyurea (C) or with or without 0.008% methyl methanesulfonate (D). Note that the full ranges of galactose concentrations were independently plated as loading/growth controls for both C and D.

